# Phage-Nanoparticle Cocktails as a Novel Antibacterial Approach: Synergistic Effects of Bacteriophages and Green-Synthesized Silver Nanoparticles

**DOI:** 10.1101/2025.03.13.643104

**Authors:** Mateusz Wdowiak, Sada Raza, Mateusz Grotek, Rafał Zbonikowski, Julita Nowakowska, Maria Doligalska, Ningjing Cai, Zhi Luo, Jan Paczesny

**Affiliations:** Institute of Physical Chemistry, Polish Academy of Sciences, Marcina Kasprzaka 44/52, 01-224 Warsaw, Poland; Military University of Technology, gen. Sylwestra Kaliskiego 2, 00-908 Warsaw, Poland; University of Warsaw, Faculty of Biology, Ilii Miecznikowa 1, 02‐096 Warsaw, Poland; Laboratory of Bioinspired Medicine and Materials, Southern University of Science and Technology, 1088 Xueyuan Avenue, Shenzhen 518055, P.R. China

## Abstract

Bacteriophages have emerged as promising natural antibacterial agents, offering a targeted approach to combating bacterial infections. While phage-antibiotic cocktails are widely explored to enhance antibacterial efficacy and prevent resistance, research on phage-nanoparticle combinations remains limited. However, antibiotic resistance continues to rise, necessitating alternative strategies. Combining bacteriophages with nanoparticles presents a novel approach that could enhance antibacterial potency while reducing the risk of resistance, yet studies in this area are still scarce. We explore the synergy between green tea extract-capped silver nanoparticles (G-TeaNPs) and bacteriophages in combating pathogenic bacteria (*Staphylococcus aureus, Salmonella enterica*). G-TeaNPs show no antiphage activity, ensuring compatibility in phage-NP formulations. These combinations significantly reduce bacterial counts in a short time (only 3 hours), e.g., *S. aureus* survival is around 30% after incubations with just 0.001 mg/mL of G-TeaNPs, with G-TeaNPs and phages alone result in around 80% and 70% survival, respectively. Cytotoxicity tests against eukaryotic 3T3 NIH fibroblast cells confirmed biocompatibility at effective concentrations. Additionally, we examine G-TeaNPs’ impact on the free-living protist *Acanthamoeba castellanii*. Both green tea extract and G-TeaNPs can reduce *A. castellanii* cell counts by 80%, but only at concentrations larger than 10 mg/mL. Microscopy revealed nanoparticle uptake by amoebae, causing intracellular accumulation and vacuolization, while green tea extract induced similar changes without uptake. Our findings highlight G-TeaNPs as safe, effective agents in phage-nanoparticle antibacterial formulations with dual antimicrobial and amoebicidal properties for therapeutic and environmental applications.

## Introduction

Phages eliminate approximately 40% of bacterial biomass daily [1]. Studies on the effects of phages on cohabiting microorganisms remain rare and undervalued, even in the most complex ecosystems [2]. The most studied environments, including food processing, human guts, and plant crops, have much to learn regarding their environmental phages and impact in various contexts [1].

Though control of bacterial pathogens by lytic phages (phage therapy) has been around for almost a century, the recent emergence of antibiotic-resistant bacteria has led to renewed interest in phage research. The therapeutic, antibacterial application of phages became known as phage therapy, especially in clinical or veterinary contexts [3]. More broadly, phages have also been used as biological control agents, reducing bacterial loads in foods, such as *Listeria monocytogenes* in food processing, zoonotic pathogens in food animals, or treating crops against plant pathogenic bacteria [4]. Since the 21st century, more studies have enhanced the understanding of phage biology and immunology, ensuring the safety of phage therapy with modern technologies like whole-genome sequencing and automation [5]. Phage therapy is currently used to treat various infections, and the FDA has approved clinical trials indicating no safety concerns. Renewed research and trials have shown phage therapy’s efficacy and safety, addressing the growing threat of antibiotic-resistant infections [6].

Phages alone have been utilized in several applications, such as the ‘*Phagoburn*’ project, which aimed to treat burn wound infections [7]. However, the main disadvantage relates to identifying appropriate lytic phages with high virulence and a broad spectrum of bacterial hosts to suit different patients [8]. The commercialization of phage preparations is infrequent due to high costs and patenting difficulties [9]. Regulatory approval for large-scale use is also lacking, as seen in Phagoburn, where improper storage conditions dramatically decreased phage titers [10]. Despite these challenges, phage-based treatments are still considered potential alternatives, especially given the rise in multidrug-resistant infections.

Despite its potential, phage therapy has not received widespread regulatory approval for human use. Nanotechnology presents a powerful tool for overcoming many barriers associated with phage preparations. Nanotechnology-based advances, such as nano-encapsulation and nanofibers, offer solutions to pharmacological and clinical challenges by protecting phages from the host’s immune system and environmental factors, ensuring sustained release, and improving delivery [11]. Encapsulation can form nanovesicles that shield phages and provide controlled release [12]. Nanofibers produced through electro-spinning offer another method to enhance phage therapy outcomes [11]. Integrating nanotechnology with phage therapy optimizes the pharmacokinetic profile of phages, ensuring better therapeutic efficacy and overcoming the limitations of phage-only formulations.

Phages are often tested as cocktails with antibiotics [13]. By combining the precision of bacteriophages, which selectively target and lyse specific bacterial strains, with the broad-spectrum efficacy of antibiotics, these cocktails enhance bacterial eradication while reducing the likelihood of resistance development. Phages can disrupt bacterial biofilms, increasing antibiotic penetration and effectiveness [14], while antibiotics can weaken bacterial defenses, making them more susceptible to phage attacks [15]. There are also attempts to mix phages with other adjuvants, e.g., essential oils [16]. However, not many reports show the simultaneous action of phages and nanoparticles. For example, a combination of silver nanoparticles (AgNPs) and bacteriophage ZCSE2 demonstrated enhanced antibacterial properties against *Salmonella*, with effective MIC and MBC values at 23 μg/mL [17]. Similarly, gold nanorods conjugated with chimeric M13 phages, forming “*phanorods*", effectively targeted *P. aeruginosa* biofilms in wound infections, highlighting their potential for treating superficial infections [18]. Another study synthesized Ag-CS nanoparticles with phages, achieving significant bactericidal and inhibitory effects against gram-positive and gram-negative bacteria and biofilm production with MIC values between 31.2 and 62.2 μg/mL and MBC values between 15.6 and 500 μg/mL [19]. Additionally, T7 phages armed with silver nanoparticles efficiently eradicated bacterial biofilms and were non-toxic to eukaryotic cells, indicating a synergistic and safe strategy for biofilm treatment [20]. Beyond bacterial infections, the integration of therapeutic viruses and nanomaterials has also revolutionized cancer therapy, enabled precision targeting, and reduced side effects, thus presenting a novel approach to overcoming the limitations of traditional treatments [21]. These examples underscore the potential of phage-nanoparticle combinations as potent antimicrobial and therapeutic agents.

In this study, we verified if metal NPs could be effectively used against viruses. We confirmed that AgNPs, including NPs synthesized using green tea extracts, had no significant ‘*antiphagant*’ activity towards examined viruses. This observation allowed us to prepare phage-NP cocktails of superior activity toward bacterial pathogens. These formulations presented improved antibacterial activity compared to frequently used phage-antibiotics cocktails. Finally, using mammalian cells and the free-living unicellular organism *A. castellanii*, we proved that our nanoparticles were safe in dedicated concentrations for both humans and aquatic protists. Additionally, we explained the potential amoebicidal activity of green tea extracts and G-TeaNPs.

## Materials and methods

### Synthesis of C-AgNPs and G-TeaNPs

All commercially available reagents were used as received without further purification. According to the product specifications provided by the producer (Sigma Aldrich), the purity of reagents was >98%, estimated using GC analysis or titration with appropriate reagents. A control batch of silver nanoparticles was prepared following a modification of the method described by Agnihotri *et al*. [22]. An aqueous solution (48 mL) containing 2 mM of NaBH_4_ and 2 mM trisodium citrate (TSC) was heated to 60°C and stirred for 30 minutes. Next, the AgNO_3_ solution (2 mL, 11.7 mM) was added dropwise. The mixture was heated to 90°C, and the pH was adjusted to 10.5 using 0.1 M NaOH. Finally, the reaction was stirred further at 90°C for 20 minutes until an evident color change was observed.

Green tea silver nanoparticles (G-TeaNPs) were synthesized using a slightly modified method by Nakhjavani et al. [23]. Green tea leaves (Loyd, Mokate, Poland) were frozen with liquid nitrogen and ground into a fine powder using a mortar and pestle. To prepare the extract, 10 g of the ground tea leaves were brewed in 100 mL of hot distilled water (60°C) for 15 minutes. The extract was then cooled to room temperature, centrifuged twice (9000 rpm for 10 minutes, followed by 15000 rpm for 10 minutes) to remove debris, and filtered using a 0.22 μm syringe filter. For nanoparticle synthesis, 25 mL of the filtered extract was stirred in an Erlenmeyer flask while 750 μL of 10 mM AgNO3 was added dropwise. The reaction mixture was stirred for 2 hours, during which the solution’s color turned yellow-brown with a silver shine, indicating nanoparticle formation. The resulting suspension was centrifuged at 10 000 rpm for 10 minutes, and the pellet was re-dispersed in distilled water. This purification step was repeated six times. Finally, the nanoparticles were suspended in water at a concentration of 1 mg/mL and stored at 4°C for later use [24].

### DLS, zeta potential

The hydrodynamic sizes of the NPs were determined using the dynamic light scattering (DLS) technique. The DLS measurements were conducted with the Malvern ZetaSizer Nano-ZS instrument using 1 cm quartz cuvettes (Hellma, Germany).

### Scanning Electron Microscopy (SEM) and EDX

Scanning electron microscope (SEM) images were collected with a Nova NanoSEM 450 under a high vacuum (10^−7^ mbar). The purified samples were deposited on a silicon substrate, allowed to dry, and mounted onto the standard SEM stub with carbon tape. The images were collected using the Through Lens Detector (TLD) of secondary electrons at a primary beam energy of 10 kV and a 4.8 mm working distance. The average diameters of the NPs were calculated from the SEM images using the ImageJ software by measuring at least 100 particles per sample.

### Fourier-transformed infrared spectroscopy (FTIR)

FTIR studies were performed with a Vertex 80 V FTIR spectrometer (Bruker, USA) equipped with a Platinum ATR (Bruker, USA) module. The tea extracts were dried by rotary evaporation at 65°C to remove water. The G-TeaNPs were centrifuged to remove water and then dried in the oven at 65°C. Dried powder samples were placed on a diamond prism (1 mm × 1 mm) to cover it entirely. The spectral resolution of the measurement was 2 cm^-1^, and the number of scans was 64.

### PXRD

X-ray diffraction analysis was performed with a Malvern PANalytical Empyrean range diffractometer at room temperature with the wavelength λ = 0.154 nm.

### The effects of TeaNPs on bacteriophages

Three different phages were selected: T4, representing one of the most abundant phages; PAO1, known for its relatively resistant nature; and Phi6, a model for highly resistant eukaryotic viruses purchased from Phage Consultants, Poland, (T4), and DSMZ (PAO1 and Phi6, respectively). The phages were exposed to citrate-capped silver nanoparticles (C-NPs) as a control, as well as silver nanoparticles synthesized using black tea (B-TeaNPs), green tea (G-TeaNPs), and red tea (R-TeaNPs). B-TeaNPs and R-TeaNPs were synthesized similarly to G-TeaNPs [24].

The change in PFU/mL was recorded after 24 hours using a double-layer agar assay. Petri plates were first filled with 20 mL of LB agar medium and left to solidify. 4 mL of top LB agar (prepared with liquid medium and 0.5% agar instead of 1.5% agar) was cooled to around 50°C and mixed with 200 μL of the refreshed culture of the appropriate bacteria strain, and poured onto the plate. Dilutions of the phage samples were prepared, and from each dilution, 7 to 8 droplets of 5 μL solution were placed onto the top agar layer. The number of plaques was counted after incubation of the plates at 37°C for 24 h.

### Bacteriophage-AgNPs cocktails against bacterial pathogens

The bacteria were cultured according to the plating method protocol. The single colony from the culture on the LB agar plate was inoculated into 10 mL of LB and then incubated at 37°C in an orbital shaker (220 RPM) overnight (about 16 hours). Next, the overnight cultures were refreshed by adding 7.5 mL of fresh LB medium to 2.5 mL of overnight culture and incubated for one hour at 37°C. Then, the refreshed cultures were both diluted with LB to reach an optical density of around OD_600_ = 1.0 for *Salmonella enterica* DSM 18552 (about 8.0 × 10^8^ CFU/mL) (colony-forming units per mL), OD_600_ = 1.0 for *Staphylococcus aureus* ATCC 43300 (about 5.0 × 10^9^ CFU/mL). Afterward, the bacteria were suspended in a PBS buffer solution to reach the initial concentration of 10^5^ CFU/mL. Then, bacteria were exposed to a rate of infections (ROI) of adequate phages (P22, vB_SauS_CS1) of 1, 10, and 100, and concentrations of G-TeaNPs ranging from 0.1 mg/mL to 0.0001 mg/mL. Both P22 and vB_SauS_CS1 were purchased from DSMZ.

The survival rate was evaluated after only 3 h of incubation at standard conditions (220 RPM, 37°C). After the incubation, 100 mL of each sample was placed onto the fresh LB agar plates. The plates were incubated overnight at 37 °C. Then, the number of bacteria was calculated based on the colony number, according to the equation CFU per mL = N × D × 10 (N – number of colonies; D – dilution).

### Amoebae – toxicity and TEM imaging

The cytotoxic effect of reagents such as AgNPs, G-TeaNPs, and green tea extract was tested in vitro on strain 309 of *Acanthamoeba castellanii* T4 genotype (isolated from the environment [25]). Amoeba strains were cultured axenically on Bacto-Casitone liquid medium + horse serum as described by Červ [26]. Amoeba cultures were established on polystyrene plates (Nest Biotechnology Co., Ltd, Non-Pyrogenic) with 24 wells. 1 mL of cell suspension was added to the wells, and 1 mL of reagent containing a concentration of AgNPs, green tea extract, and G-TeaNPs to achieve the target dilution. Reagents were added to the axenic amoeba culture (5×10^4^ cells/mL) in the following concentrations: 0.05, 0.1, 5, 10, and 20 mg/mL. The increase or decrease in the number of amoebae was checked at 48-hour intervals using a Thom’s haemocytometer chamber. The control group was a culture of amoebae without reagents [27]. The minimum concentration of reagents that inhibited the growth of the amoeba culture was determined. The test was performed in 3 replicates, up to 7 days, of cultures stored at 28°C in a humid and dark chamber.

### Transmission electron microscope (TEM)

The cells were fixed in 2,5 % glutaraldehyde in 0,1M cacodylate buffer pH=7,2 overnight at room temperature. Samples were washed in cacodylate buffer thrice and stained with 1 % osmium tetroxide in ddH_2_O overnight at room temperature. Samples were washed in ddH_2_O and dehydrated through a graded series of ethanol (30%, 50%, 70%, 80%, 96%, absolute ethanol and acetone). Samples were embedded in epon resin (SERVA) and polymerized for 24h at 60°C in an incubator (Agar Scientific, England). Next, 70 nm sections were cut with a diamond knife on RMC MTXL ultramicrotome (RMC Boeckeler Instruments, USA). The sections on copper grids were not contrasted. Samples were analyzed in a LIBRA 120 transmission electron microscope produced by Carl Zeiss (Oberkochen, Germany) at 120 keV. Photographs were made with a Slow-Scan CCD camera (ProScan, Germany) using the EsiVision Pro 3.2 software.

### Cytotoxicity towards mammalian cells

The biocompatibility of modified gold nanoparticles was assessed using the Alamar Blue assay using 3T3 NIH fibroblast cells. All reagents were obtained from commercial suppliers: AlamarBlue reagent (ThermoFisher Scientific), Triton X-100 (Sigma-Aldrich), and Trypan Blue (ThermoFisher Scientific). 3T3 NIH fibroblast cells were cultured in Dulbecco’s Modified Eagle Medium (DMEM) supplemented with 10% fetal bovine serum (FBS) and 1% penicillin-streptomycin, in a humidified atmosphere at 37°C and 5% CO_2_. Cells were subcultured upon reaching 70–80% confluency and harvested for experiments during the log phase of growth. Cells were harvested and counted using Trypan Blue exclusion to ensure viable cell counts. Cells were diluted to a concentration of 7.5 × 10 cells/mL, and 200 μL of cell suspension (equivalent to 1.5 × 10^4^ cells) was seeded into each well of a 96-well tissue culture-treated plate. The cells were incubated overnight at 37°C with 5% CO_2_ to allow for surface attachment. Cells were treated with nanoparticles at a 0.1 mg/mL concentration the following day. The cells were incubated with the nanoparticles for 24 and 48 hours at 37°C in a humidified atmosphere with 5% CO_2_. After each incubation period, the medium was aspirated and replaced with fresh medium containing 10% (v/v) Alamar Blue solution. The cells were incubated with the AlamarBlue mixture for 4 hours at 37°C with 5% CO_2_. After incubation, 150 μL of the medium from each well was transferred to a new 96-well plate, and fluorescence was measured using a plate reader with excitation at 530–570 nm and emission at 580–620 nm. The blank value (from wells without cells) was subtracted from each reading to ensure accuracy. Cell viability was calculated based on metabolic activity, which was directly proportional to the Alamar Blue fluorescence intensity.

Cytotoxicity tests were also performed using MTT proliferation/metabolic activity assays. We used a cancer cell line - HeLa (cervical cancer). The MTT assay was performed using around 10,000 cells/well for both tested cell lines (controlled with Countess II Cell Counter). Cells were seeded into a 96-well plate (Greiner Bio-One) and incubated in an incubator for 24 hours at 37°C. Then, the medium was removed, and the tested formula (containing dye and polymer) at five different concentrations (double dilutions) was added to the cell fresh medium. The experiments were repeated five times for each concentration. After 6 hours of incubation, the medium was replaced with a culture medium including 1 mM 3-(4,5-dimethylthiazol-2-yl)-2,5-diphenyltetrazolium bromide (MTT reagent, Thermo Fischer Scientific). Cells were incubated with MTT reagent for 3 hours at 37°C. Then, solutions were replaced with DMSO and incubated for 10 minutes. The absorbance in each well was measured at 540 nm using a SpectraMax i3x MultiMode Microplate Reader with injectors (Molecular Devices). For each experiment, negative and positive controls were included. Negative controls contained 1% Triton-X 100, while positive controls were cells cultured under standard conditions without a tested formula.

## Results

### Characterization of G-TeaNPs

The SEM image of the silver nanoparticles (AgNPs) capped with green tea extract (G-TeaNPs) revealed their morphology, with agglomerates bound by a thin layer likely originating from the tea extract coating (**Figure 1a**). The size distribution of the nanoparticles was analyzed using ImageJ (**Figure 1b**), showing that most particles fell within the 40–50 nm range. However, a noticeable deviation from a normal distribution was observed due to an overrepresentation of particles around ∼20 nm. The average particle diameter was calculated to be 45.6 ± 15.6 nm. This irregular size distribution is consistent with findings from our previous study, where G-TeaNPs’ efficacy against bacteria was evaluated [24].

**Figure 1.**
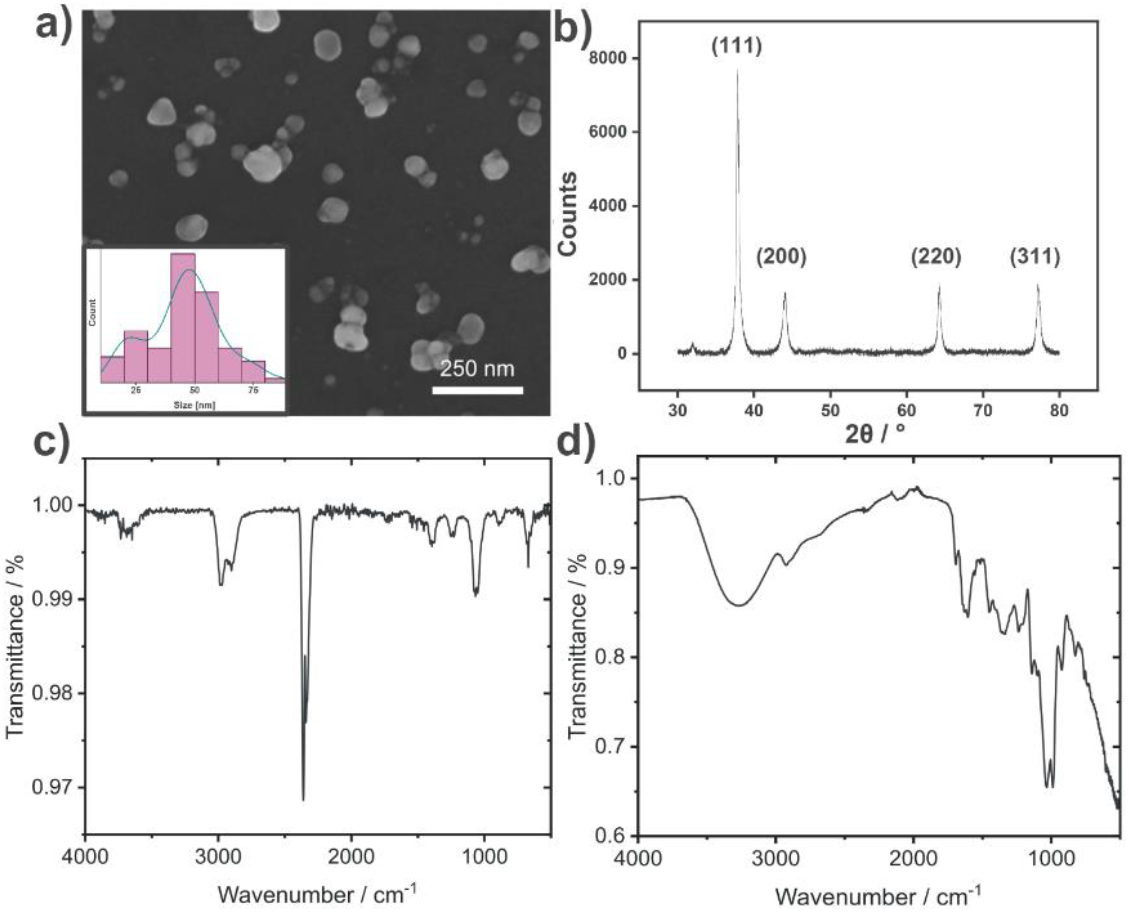
**a)** SEM image of G-TeaNPs (silver nanoparticles synthesized using green tea extract), inset: size distribution (diameter), line - kernel density. **b)** Diffractogram of AgNPs capped with green tea extract. **c)** FTIR spectra of G-TeaNPs and **(d)** green tea extract.

The nanoparticles were further analyzed using Fourier-transform infrared spectroscopy (FTIR), revealing spectral features consistent with those reported in our previous study (**Figure 1** and **1d**). A broad band in the range of 3000–3600 cm−^1^, characteristic of O–H stretching, was observed. This band is typical of alcohols, phenols, and carboxylic acids commonly found in tea extract components, as well as possibly due to adsorbed water molecules. Additional prominent bands included C–H stretching (2850–2950 cm−^1^), C=O stretching (1700–1750 cm−^1^), C=C stretching (1620–1680 cm−^1^), N–H bending (1500–1600 cm−^1^), and C–O stretching (1050–1300 cm−^1^). These bands corresponded to functional groups commonly found in tea extract constituents, such as catechins, flavonoids, phenolic acids, and amino acids. Notably, except for the broad O–H stretching band, the same characteristic peaks were observed in the FTIR spectrum of the G-TeaNPs, confirming the successful capping of nanoparticles by the tea extract components.

Energy-dispersive X-ray spectroscopy (EDX) was performed on the AgNPs sample, with the Si substrate used as a reference background. The analysis confirmed the presence of silver in the sample (**Figure S1**).

The X-ray diffraction (XRD) analysis of the sample displayed a characteristic diffractogram of AgNPs, i.e., the face-centered cubic crystalline structure of metallic silver (JCPDS 01-071-4613) (**Figure 1b**). Using Scherrer’s equation, the crystallite size was calculated from the four prominent diffraction peaks (with a shape factor of 0.94), yielding an average size of 19.8 ± 3.4 nm. This suggests that a significant proportion of the nanoparticles are not single crystals, which may also account for the overrepresentation of ∼20 nm particles observed in the size distribution analysis.

We also analyzed two peaks at 32.0° and 35.9° of low intensity. They were associated with the highly intense peaks of silver oxide species, such as Ag_2_O, AgO, and Ag_2_O_3_ [28]. These findings indicated that the oxidation of the silver nanoparticles during synthesis or storage was minimal.

### Antimicrobial activity of G-TeaNPs

Before testing phage-nanoparticle cocktails against pathogenic bacteria, evaluating the potential effects of the nanoparticles (NPs) on phages is essential. **Figure S2** confirmed that our nanoparticles exhibited no significant inactivation effects on the tested phages. While PAO1 showed a minor reduction of approximately 0.5 log compared to the control group containing only phages, there was no observable difference between the effects of citrate-capped nanoparticles and tea-based nanoparticles on PAO1. This suggested that any slight reduction in PFU/mL was not attributable to the functionalization of the tea extract. For T4 and Phi6, no measurable decrease in phage activity was observed for any of the tested nanoparticles, indicating the lack of antiviral activity of AgNPs, including the G-TeaNPs, against both enveloped and non-enveloped viruses. These results indicated that the nanoparticles, whether citrate-capped or tea-capped, did not possess phage-inactivating properties, supporting their compatibility for further testing in phage-nanoparticle combinations targeting pathogenic bacteria.

Once it was established that the nanoparticles did not have any direct inactivating effect on phages, we proceeded to test phage-nanoparticle (phage-NP) cocktails using green tea silver nanoparticles. G-TeaNPs were previously identified as the most effective against bacteria and fungi among a series of silver particles prepared with extracts from various teas [24].

For the novel phage-NPs cocktails, we selected *S. enterica*, one of the most frequently occurring foodborne pathogens (a gram-negative bacterium), and a methicillin-restraint strain of *S. aureus* (MRSA; a gram-positive bacterium). The efficacy of these combinations was assessed across a range of rates of infection (ROI, representing the ratio of phages to bacterial cells). In **Figure 2a**, ROI = 1 was kept constant, but the concentration of G-TeaNPs varied from 0.1 mg/mL to 0.0001 mg/mL. **Figure 2b** shows the results of experiments where the concentration of G-TeaNPs was constant (0.001 mg/mL), but ROIs varied from 1 to 100.

**Figure 2.**
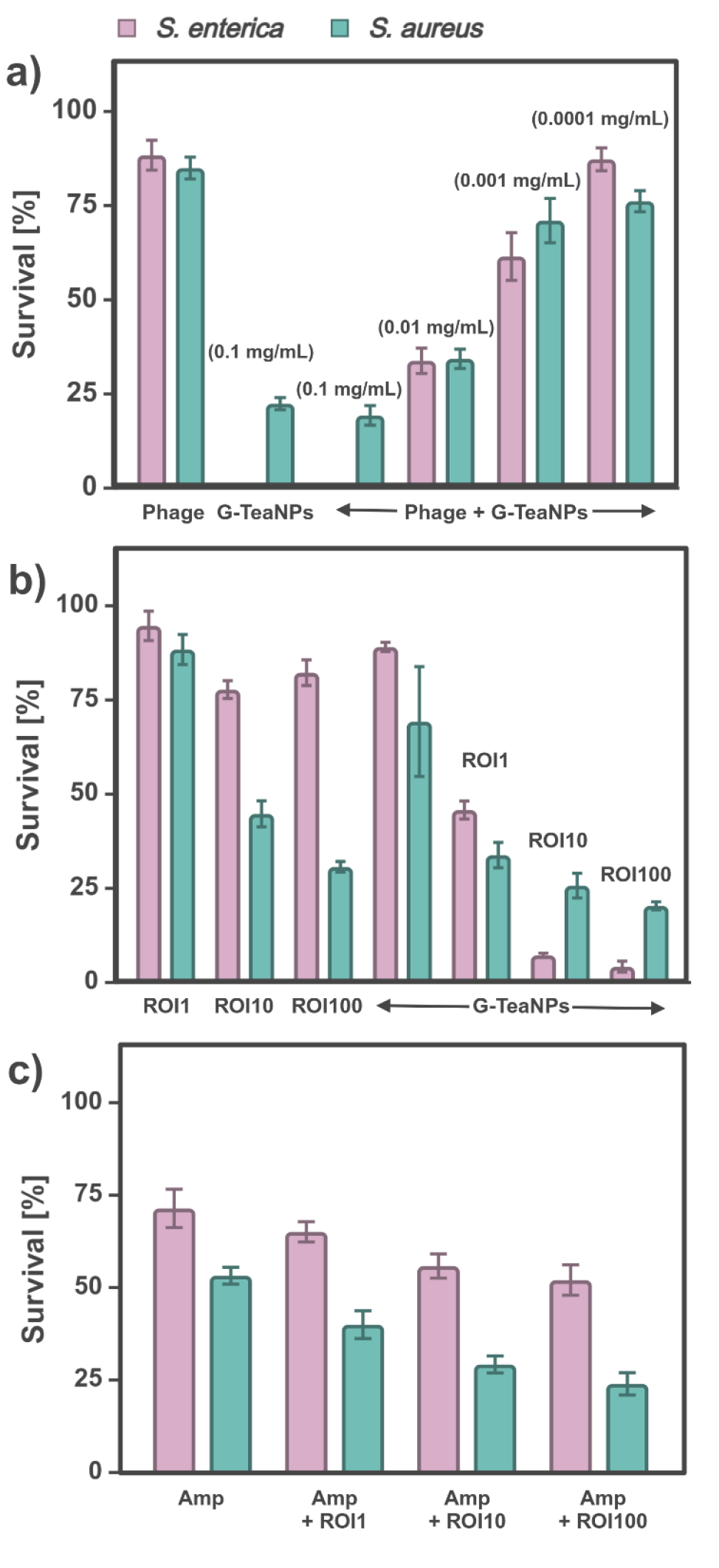
Antibacterial activity of phage-nanoparticle (phage-NP) cocktails and phage-antibiotic combinations against *Staphylococcus aureus* (gram-positive) and *Salmonella enterica* (gram-negative). **(a)** Bacterial survival after treatment with G-TeaNPs (0.1 mg/mL to 0.0001 mg/mL) combined with P22 (*S. enterica*) or vB_SauS_CS1 (*S. aureus*) bacteriophages (ROI = 1). **(b)** Effect of phage-NP cocktails ([G-TeaNPs] = 0.001 mg/mL) at different rates of infection (ROI = 1 to 100). **(c)** Comparison of bacterial survival following treatment with ampicillin (AMP) alone or in combination with bacteriophages at varying ROIs.

Adding phages at an ROI = 10 or higher significantly enhanced antibacterial effects, allowing for a substantial reduction in the required working concentrations of G-TeaNPs. Specifically, with phages at ROI = 10, the bacterial survival rate decreased to 10–20%, depending on the species, even when the concentration of G-TeaNPs was as low as 0.001 mg/mL (1 μg/mL).

In contrast, a similar efficacy with G-TeaNPs alone required a 100 times higher concentration (0.1 mg/mL).

For comparison with commonly used phage-antibiotic cocktails, we tested the combined effects of bacteriophages and ampicillin (AMP) against *S. aureus* and *S. enterica* across various rates of infection (ROIs). The results showed that the AMP-phage combination was more effective against *S. aureus* (**Figure 2d**). Specifically, AMP alone reduced bacterial survival to approximately 60%, while adding phages further reduced survival to around 30% at ROI 10 and 25% at ROI 100.

In the case of *S. enterica*, the impact of the AMP-phage combination was less pronounced but still significant. AMP alone decreased bacterial survival to approximately 75%. Adding phages reduced survival to around 70% (AMP + ROI 1) and 58% (AMP + ROI 100). Notably, the phage-NP cocktails at ROIs 10 and 100 achieved far greater reductions in survival, with bacterial survival rates dropping to approximately 10%.

The overall efficacy of the AMP-phage combination was similar to or better than that of the phage-NP cocktails. This might be crucial when fighting multidrug-resistant bacteria, where antibiotics are inefficient and novel agents are needed. We also underline that these results were obtained after only 3 h incubation.

### Cytotoxicity towards amoebae and mammalian cells

To assess potential environmental impacts and effects on free-living unicellular organisms, *Acanthamoeba castellanii* was used as a model organism. Amoebae were exposed to tea extracts, citrate-capped AgNPs, and G-TeaNPs for seven days, with live amoebae counts taken every two days (**Figure 3d**). The experiments revealed that G-TeaNPs were ineffective against *A. castellanii* at low concentrations. Depending on the batch of particles, up to 5 mg/mL did not affect amoebae. In the case of these particles, the lowest concentration inhibiting the growth was 10 mg/mL, a dose approximately 16 times higher than the cytotoxic threshold for mammalian cells. Interestingly, no clear correlation was observed between G-TeaNP concentration and the formation of amoebae cysts. Surprisingly, green tea extracts alone were as effective in inhibiting amoebae growth as G-TeaNPs, a phenomenon not observed in antibacterial assays.

**Figure 3.**
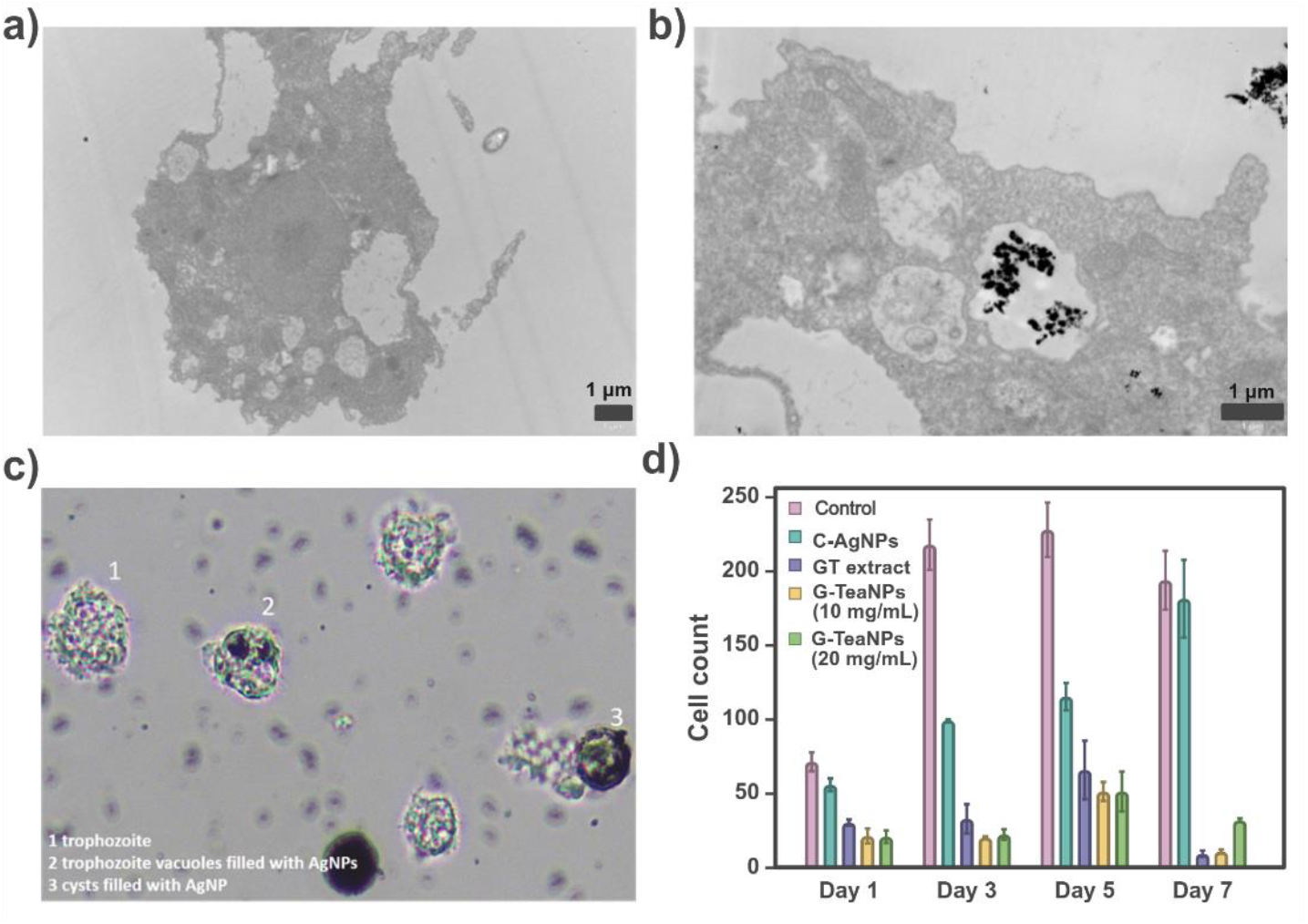
**b)** TEM images of *A. castellanii* exposed to G-TeaNPs, showing nanoparticle uptake and accumulation within intracellular vacuoles, compared to control cells **(a). c)** Cells exposed to G-TeaNPs exhibit increased vacuolization and morphological changes: 1) trophozoites, 2) trophozoite vacuoles filled with AgNPs, and 3) cysts containing AgNPs. Similar vacuolization and size/shape reduction were observed in cells treated with green tea extract but without nanoparticle accumulation. These findings highlight the cellular uptake of nanoparticles and their potential influence on *A. castellanii* morphology. **d)** Cytotoxic effects of citrate-capped C-AgNPs, G-TeaNPs, and green tea extract (GT) on *Acanthamoeba castellanii* over 7 days. G-TeaNPs inhibited amoebae growth only at high concentrations, while lower concentrations (i.e., 0.1 mg/mL as used in the other assays) were ineffective. GT alone was as effective as G-TeaNPs in reducing amoebae counts.

Transmission electron microscopy confirmed that *A. castellanii* actively phagocytosed the nanoparticles, accumulating within intracellular vacuoles (**Figure 3b**). Light microscopy further revealed distinct morphological changes in both live amoebae and cysts after exposure to G-TeaNPs (**Figure 3c**). Specifically, trophozoites and cysts were visibly filled with AgNPs within their vacuoles after five days of exposure. Exposure to green tea extract alone also induced noticeable reductions in cell size and shape and increased vacuolization, similar to what was observed in cells treated with G-TeaNPs. However, no nanoparticle accumulation was observed in the case of green tea extract.

The control sample showed normal *A. castellanii* forms, including moving trophozoites, resting trophozoites, and cysts, while cells exposed to G-TeaNPs display increased vacuolization, with AgNPs accumulating inside trophozoites and cysts. Similar vacuolization and morphological changes were observed in cells treated with green tea extract, although without the visible presence of nanoparticles.

These findings highlight the cellular uptake of nanoparticles by *A. castellanii* and their potential influence on amoebae morphology. Additionally, the inhibitory effect of green tea extract on *A. castellanii* raises questions about its mechanism of action, independent of nanoparticles, and its broader implications for environmental and biological systems. Importantly, these results also underscore the species-specific effects of G-TeaNPs, with significant differences in cytotoxicity between mammalian cells and amoebae.

To evaluate the biocompatibility of G-TeaNPs nanoparticles, an Alamar Blue assay was conducted using the 3T3 NIH fibroblast cell line. Two types of nanoparticles were tested: citrate-capped silver nanoparticles (AgNPs) and G-TeaNPs. The chosen concentrations were 1 mg/mL and 0.1 mg/mL, based on their effective antibacterial activity.

3T3 NIH cells were treated with different nanoparticle formulations, including AgNPs and G-TeaNPs, for 24 hours. Cell viability was assessed using the Alamar Blue assay, which measured the conversion of resazurin to its fluorescent product, resorufin. The quantitative results in **Figure 4f** demonstrate that cell viability exceeded 80% for G-TeaNPs at 0.1 mg/mL, while that of AgNPs remained below 60%. At 1 mg/mL, both the nanoparticles showed cell viability below 5%. Observations of cell morphology further confirmed nanoparticle biocompatibility.

**Figure 4.**
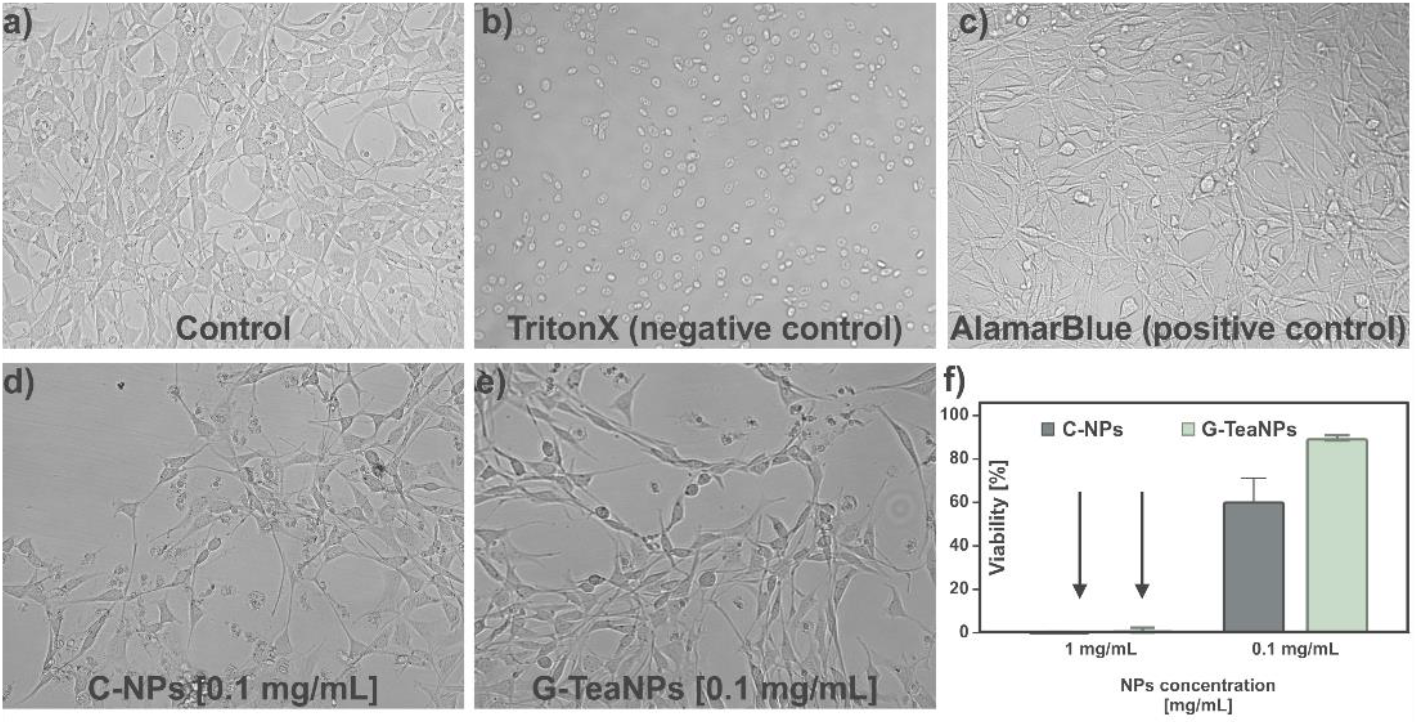
Optical microscopic images of 3T3 NIH fibroblast cells after 24 hours of incubation with different nanoparticle treatments: **a)** Untreated control cells, **b)** Cells treated with 1% TritonX as a negative control. **c)** Cells treated with only AlamarBlue (positive control). **d)** Cells treated with control silver nanoparticles (C-AgNPs, 0.1 mg/mL), **d)** Cells treated with G-TeaNPs (0.1 mg/mL), **e)** Viability of cells treated with AgNPs, and G-TeaNPs after 24 hours of incubation at 1 mg/mL and 0.1 mg/mL.

Untreated control cells (**Figure 4a**) maintained their characteristic elongated, spindle-like morphology, typical of healthy 3T3 fibroblasts. Similarly, cells exposed to AlamarBlue as a positive control (**Figure 4c**) maintained their morphology. In contrast, cells treated with Triton X-100 (1%; **Figure 4b**), i.e., negative control, showed spherical, shrunken morphologies indicative of cell death, confirming the assay’s sensitivity in detecting cytotoxic effects. Cells in the presence of AgNPs and GTNPs displayed no significant morphological changes compared to controls at 0.1 mg/mL (**Figure 4 d-e**). These results suggest that G-TeaNPs retain high biocompatibility with 3T3 NIH cells, with cell viability consistently above 80%.

The cytotoxicity of citrate-capped AgNPs and G-TeaNPs was examined using the MTT metabolic activity assay on HeLa cells, with results summarized in **Figure S3**. G-TeaNPs demonstrated significantly higher toxicity compared to citrate-capped AgNPs across all tested concentrations. At 2.5 mg/mL, G-TeaNPs reduced HeLa cell survival to approximately 20%, while C-AgNPs allowed over 50% survival. Even at lower concentrations, such as 1.25 mg/mL and 0.625 mg/mL, G-TeaNPs showed consistently greater cytotoxic effects, with survival rates dropping below 70%, compared to C-AgNPs, where survival remained above 70%. Notably, the concentration of G-TeaNPs required to reach <70% cell survival (0.625 mg/mL) was over six times higher than the working concentration used in antibacterial assays and more than 600 times higher than the amount of G-TeaNPs used in phage-nanoparticle cocktails. These findings highlight the higher cytotoxicity of G-TeaNPs to mammalian cells and the importance of optimizing doses to ensure antimicrobial efficacy while minimizing potential toxicity.

## Conclusion

This study demonstrated the potential of green tea extract-capped silver nanoparticles (G-TeaNPs) as multifunctional agents, combining antimicrobial efficacy with broad biological applicability. Detailed characterization confirmed their nanoscale morphology, irregular size distribution, and functional group composition, which were consistent with green tea extract capping. Despite minor deviations, such as partial oxidation and the presence of aluminum compounds from the tea extract, the nanoparticles were stable and well-suited for subsequent applications. G-TeaNPs did not exhibit direct phage-inactivating properties, ensuring their compatibility in phage-nanoparticle cocktails. These cocktails proved particularly effective against gram-negative bacteria, such as *S. enterica*, achieving bacterial survival rates of approximately 10% at low nanoparticle concentrations (0.001 mg/mL). This highlights their ability to enhance antibacterial performance while minimizing nanoparticle usage.

Cytotoxicity testing revealed the dose-dependent effects of G-TeaNPs on 3T3 NIH, HeLa cells, and *A. castellanii*. Although G-TeaNPs were more toxic to mammalian cells compared to citrate-capped AgNPs, the effective doses used in antibacterial assays were substantially lower than the cytotoxic thresholds. Interestingly, green tea extracts alone displayed amoebicidal effects comparable to G-TeaNPs, raising questions about their intrinsic activity and environmental implications. Microscopy studies confirmed nanoparticle uptake and accumulation in amoebae, with associated morphological changes, while green tea extract induced similar changes without nanoparticle presence. This indicates that AgNPs may act primarily as carriers of tea extract components inside amoeba cells in such systems.

Overall, the results underscore the versatility of G-TeaNPs in bacterial and phage-based applications while emphasizing the importance of dose optimization to balance efficacy with cytotoxicity. These findings provide valuable insights into the mechanisms and environmental impact of green tea-derived nanomaterials, paving the way for future studies in targeted therapeutic and ecological applications.

## Author Contributions

Conceptualization: M.W., S.R., J.P.; methodology: M.W., S.R., M.D.; validation: M.W., S.R., M.G., M.D., Z.L.; formal analysis: M.W., S.R., J.P., M.D.; investigation: M.W., S.R., M.G., R.Z., J.N., M.D., N.C.; resources: J.P., M.D., Z.L.; data curation: S.R., M.W.; writing—original draft preparation, S.R.; writing—review and editing, S.R., M.W., J.P.; visualization, S.R.; supervision, J.P.; project administration, S.R., J.P.; funding acquisition, J.P.

All authors have read and agreed to the published version of the manuscript.

## Data availability

The raw data required to reproduce these findings are available to download from https://doi.org/10.18150/YBVYR9.

---

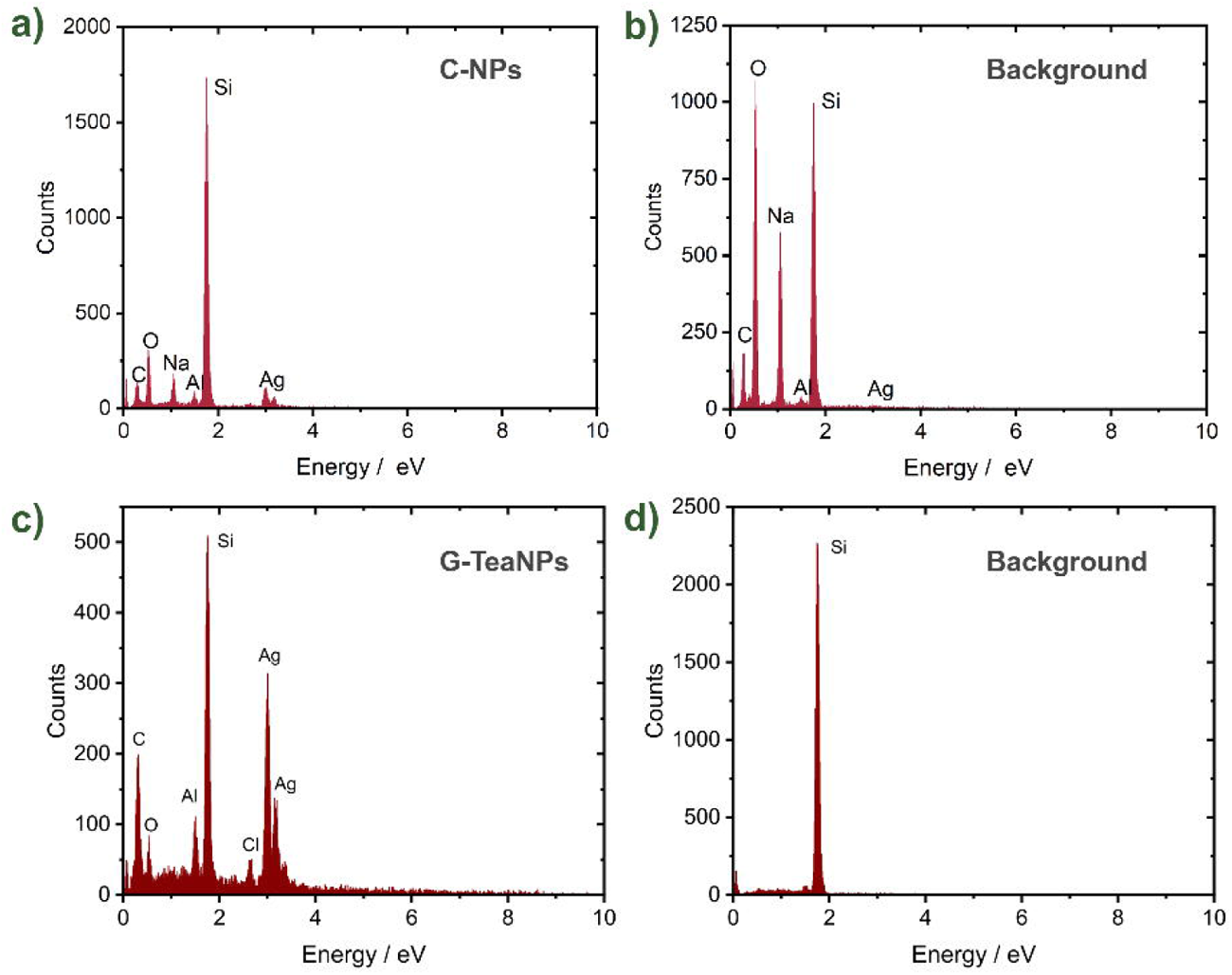

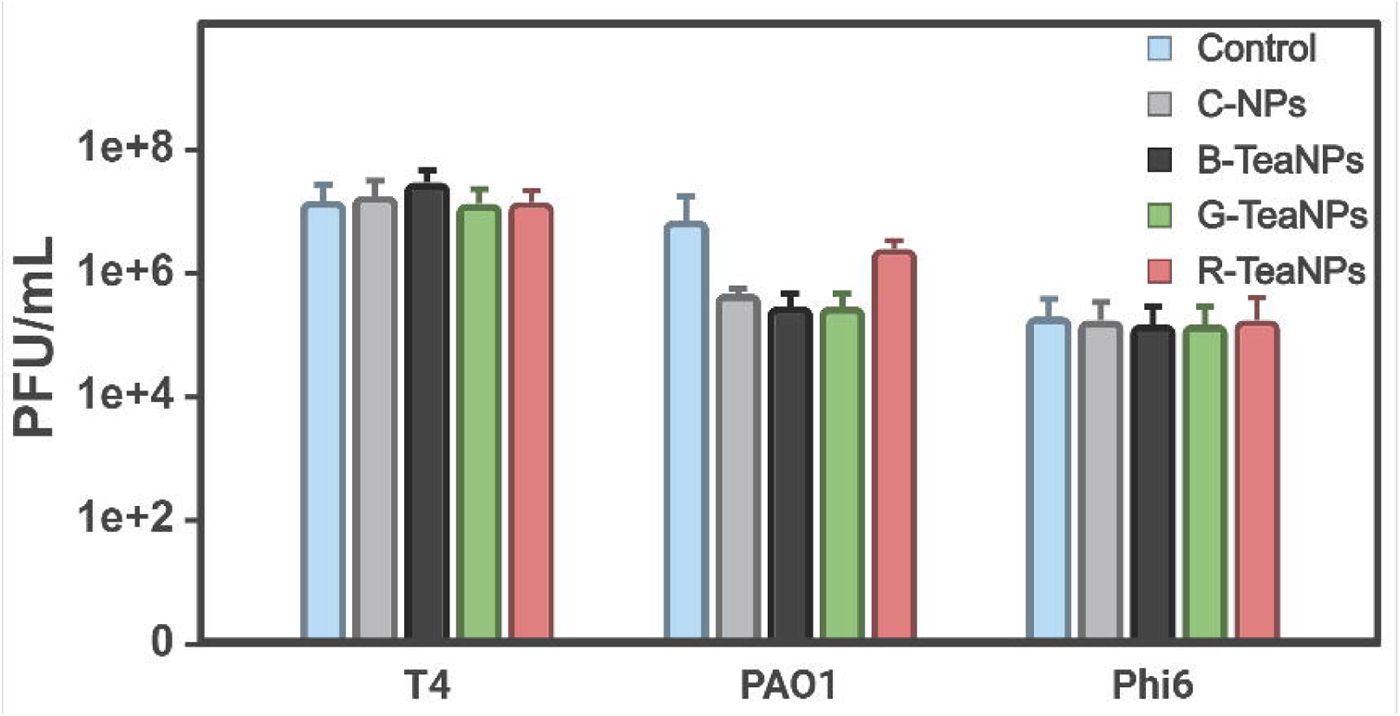

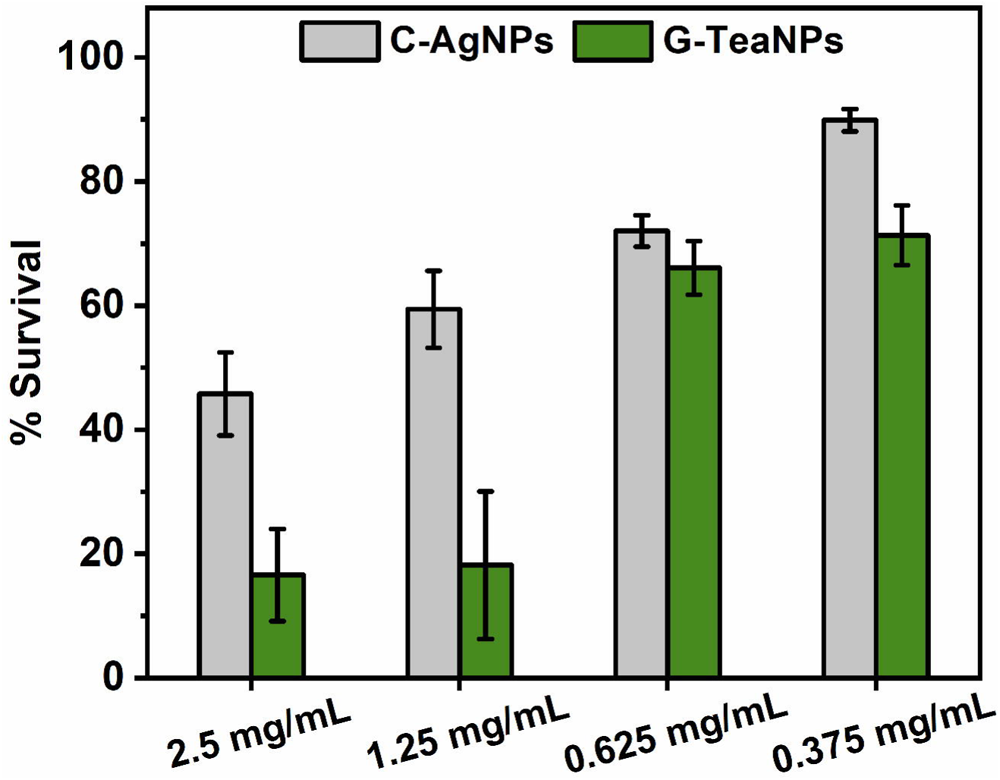

