## Supplementary Information for "Phage-Nanoparticle Cocktails as a Novel Antibacterial Approach: Synergistic Effects of Bacteriophages and Green-Synthesized Silver Nanoparticles"

Phage-Nanoparticle Cocktails as a Novel Antibacterial Approach: Synergistic Effects of Bacteriophages and Green-Synthesized Silver Nanoparticles
Mateusz Wdowiak1, Sada Raza1,*, Mateusz Grotek1,2, Rafał Zbonikowski1, Julita Nowakowska3, Maria Doligalska3*, Ningjing Cai4, Zhi Luo4, Jan Paczesny1,*

1 Institute of Physical Chemistry, Polish Academy of Sciences, Marcina  Kasprzaka 44/52, 01-224 Warsaw, Poland
2 Military University of Technology, gen. Sylwestra Kaliskiego 2, 00-908 Warsaw, Poland
3 University of Warsaw, Faculty of Biology, Ilii Miecznikowa 1, 02?096 Warsaw, Poland
4 Laboratory of Bioinspired Medicine and Materials, Southern University of Science and Technology, 1088 Xueyuan Avenue, Shenzhen 518055, P.R. China


Figure S1. Energy-dispersive X-ray spectra of the AgNPs capped with citrate (control, (a)), its background measurement (b), green tea extract (c) and its background measurement (d).

Before testing phage-nanoparticle cocktails against pathogenic bacteria, it is important to evaluate the potential effects of the nanoparticles (NPs) on phages. To this end, three different phages were selected: T4, representing one of the most abundant phages; PAO1, known for its relatively resistant nature; and Phi6, a model for highly resistant eukaryotic viruses. The phages were exposed to citrate-capped silver nanoparticles (C-NPs) as a control, as well as silver nanoparticles synthesized using black tea (B-TeaNPs), green tea (G-TeaNPs), and red tea (R-TeaNPs), following the previous research [1].
      As shown in Figure S2, none of the nanoparticles exhibited significant inactivation effects on the tested phages. While PAO1 showed a minor reduction of approximately 0.5 log compared to the control group containing only phages, there was no observable difference between the effects of citrate-capped nanoparticles and tea-based nanoparticles on PAO1. This suggests that any slight reduction in PFU/mL is not attributable to the tea extract functionalization. For T4 and Phi6, no measurable reduction in phage activity was observed for any of the tested nanoparticles, indicating the lack of antiviral activity of AgNPs, including the TeaNPs, against both enveloped and non-enveloped viruses.


Figure S2. The effect of citrate-capped silver nanoparticles (C-NPs) and tea-extract-functionalized silver nanoparticles (B-TeaNPs, G-TeaNPs, and R-TeaNPs) on the viability of three phages: T4, PAO1, and Phi6. Phage titers are presented as plaque-forming units per milliliter (PFU/mL).


Figure S3. Cytotoxicity of citrate-capped AgNPs, and G-TeaNPs examined using the MTT metabolic activity assay on HeLa cells. In all examined concentrations, G-TeaNPs presented significantly higher toxicity compared to citrate-capped AgNPs. Both types of nanoparticles were toxic to HeLa (survival <70%) starting from 0.625 mg/mL concentration. Such concentration was over 6 times higher than the working concentration of G-TeaNPs in the antibacterial assay, and over 600 times higher compared to the amount of G-TeaNPs in cocktails with phages.
